# Spatially heterogeneous structure-function coupling in haemodynamic and electromagnetic brain networks

**DOI:** 10.1101/2022.12.14.520453

**Authors:** Zhen-Qi Liu, Golia Shafiei, Sylvain Baillet, Bratislav Misic

## Abstract

The relationship between structural and functional connectivity in the brain is a key question in connectomics. Here we quantify patterns of structure-function coupling across the neocortex, by comparing structural connectivity estimated using diffusion MRI with functional connectivity estimated using both neurophysiological (MEG-based) and haemodynamic (fMRI-based) recordings. We find that structure-function coupling is heterogeneous across brain regions and frequency bands. The link between structural and functional connectivity is generally stronger in multiple MEG frequency bands compared to resting state fMRI. Structure-function coupling is greater in slower and intermediate frequency bands compared to faster frequency bands. We also find that structure-function coupling systematically follows the archetypal sensorimotor-association hierarchy, as well as patterns of laminar differentiation, peaking in granular layer IV. Finally, structure-function coupling is better explained using structure-informed inter-regional communication metrics than using structural connectivity alone. Collectively, these results place neurophysiological and haemodynamic structure-function relationships in a common frame of reference and provide a starting point for a multi-modal understanding of structure-function coupling in the brain.

## INTRODUCTION

The relationship between brain structure and function is a central concept in neuroscience [116, 138]. The complex network of synaptic projections forms a hierarchy of nested and increasingly polyfunctional neural circuits that support perception, cognition and action [10]. Modern imaging technology permits high-throughput reconstruction of neural circuits across spatiotemporal scales, and across species [76]. These comprehensive wiring diagrams of the nervous system – termed structural connectivity networks – represent the physical connections between neural elements [136]. Structural connectivity promotes electrical signaling and synchrony among distant neuronal populations, giving rise to coherent neural dynamics, measured as regional time series of electromagnetic or haemodynamic neural activity. Systematic co-activation among pairs of regions can be used to map functional connectivity networks [19, 50], which exhibit reproducible and similarly organized patterns in both task-driven and task-free paradigms [34, 107, 161]. How do we evaluate the relationship between structure and function in the human brain? The most common approach is to estimate whole-brain diffusion MRI-derived structural connectivity and functional connectivity at a desired spatial scale, and study their correspondence. Conventional models of structure-function coupling typically utilize resting-state fMRI-estimated functional connectivity, and compute a single, global cross-correlation statistic between structure and function. In these studies, the upper or lower triangle of the structural and functional connectivity matrices are vectorized and correlated with each other, revealing consistent but moderate correspondence between structural and functional connectivity (*R*^2^ *<* 0.25 in most reports) [33, 64, 65, 130] (see [138] for a review).

Two important limitations of this paradigm are increasingly recognized. First, BOLD activity only indirectly reflects the underlying patterns of electrical neural activity, mainly because of slow neurovascular coupling [9, 41, 67, 73, 129]. As a result, emerging efforts emphasize integration of multiple neurophysiological modalities, particularly magnetoenecephalography (MEG) which offers excellent temporal resolution as well as high spatial specificity when used in conjunction with source modeling [8, 115, 116]. Second, most current studies model a single, globally-uniform structure-function relationship across the brain. Recent reports, however, suggest that structure-function coupling is regionally heterogeneous, with stronger correspondence between structural and functional connectivity in uni-modal cortex, and weaker correspondence in transmodal cortex [12, 108, 151, 157, 163], potentially reflecting underlying molecular and cytoarchitectural gradients [12, 13, 48, 138, 151]. Altogether, a more detailed biological understanding of structure-function relationships – one that takes into account both neurophysiological activity and regional heterogeneity – is necessary [138].

Here we seek to comprehensively characterize region-specific patterns of structure-function coupling using neurophysiological activity. We focus on functional connectivity derived using MEG recordings, as this form of connectivity has been shown to convey richer temporal features and is overall a more veridical representation of fast neurophysiological dynamics compared to fMRI [8, 115]. Recent reports have established overall similar but only partially overlapping connectivity patterns between fMRI and MEG connectivity [23, 82, 127]. Initial studies linking structural connectivity and MEG functional connectivity also showed modest but slightly greater coupling than with fMRI connectivity, with considerable variation across canonical frequency bands [29, 91, 116, 134, 141]. Several trimodal comparisons of dMRI, fMRI, and MEG have confirmed these relationships [28, 52, 90, 116, 142]. However, why structure-function coupling is regionally heterogeneous, and how this spatial heterogeneity differs across modalities, remains unknown. Moreover, the contribution of local cytoarchitectural variation to regional patterns of structure-function coupling estimated from neurophysiological activity is unclear [18, 62, 147].

In the present report, we comprehensively benchmark the correspondence between dMRI-derived anatomical connectivity and functional connectivity across resting-state fMRI and MEG canonical frequency bands. We estimate regional patterns of structure-function coupling using a multilinear regression model that takes into account communication dynamics [125, 151]. We then explore the relationship between structure-function coupling patterns and cognitive systems, network features and cytoarchitectural profiles.

## RESULTS

The results are organized as follows. We first estimate structure-function coupling between dMRI-derived structural connectivity and a number of functional connectivity matrices, including resting-state band-limited MEG connectivity and fMRI functional connectivity. We then describe how structural-function coupling systematically varies across the neocortex from the perspective of topological, intrinsic functional, hierarchical, and cytoarchitectural organization. Structural and functional connectivity matrices were reconstructed using data from the same participants in the Human Connectome Project (HCP; dMRI, fMRI, MEG; *N* = 33 healthy young adults) [148]. All data were parcellated using the Schaefer400x7 atlas [119].

### Connectomic representations of structure and function

Fig. 1 shows the structural and functional matrices. The sparse structural connectivity matrix was estimated from dMRI (Fig. 1a). This matrix was then converted to multiple “predictor” communication matrices that represent the propensity for brain regions to communicate with each other via the structural connectivity according to different protocols. Although not an exhaustive list, we included measures that (a) have been studied before in the network neuroscience literature, and (b) were biologically plausible. Measures include Euclidean distance, shortest path length, navigation efficiency, search information, communicability and diffusion efficiency (see *Methods* for definitions). These predictors can be thought of as residing on a spectrum, from decentralized, diffusion-like communication processes (network diffusion) to centralized, routing-like communication processes (shortest path routing) [5, 14].

**Figure 1.**
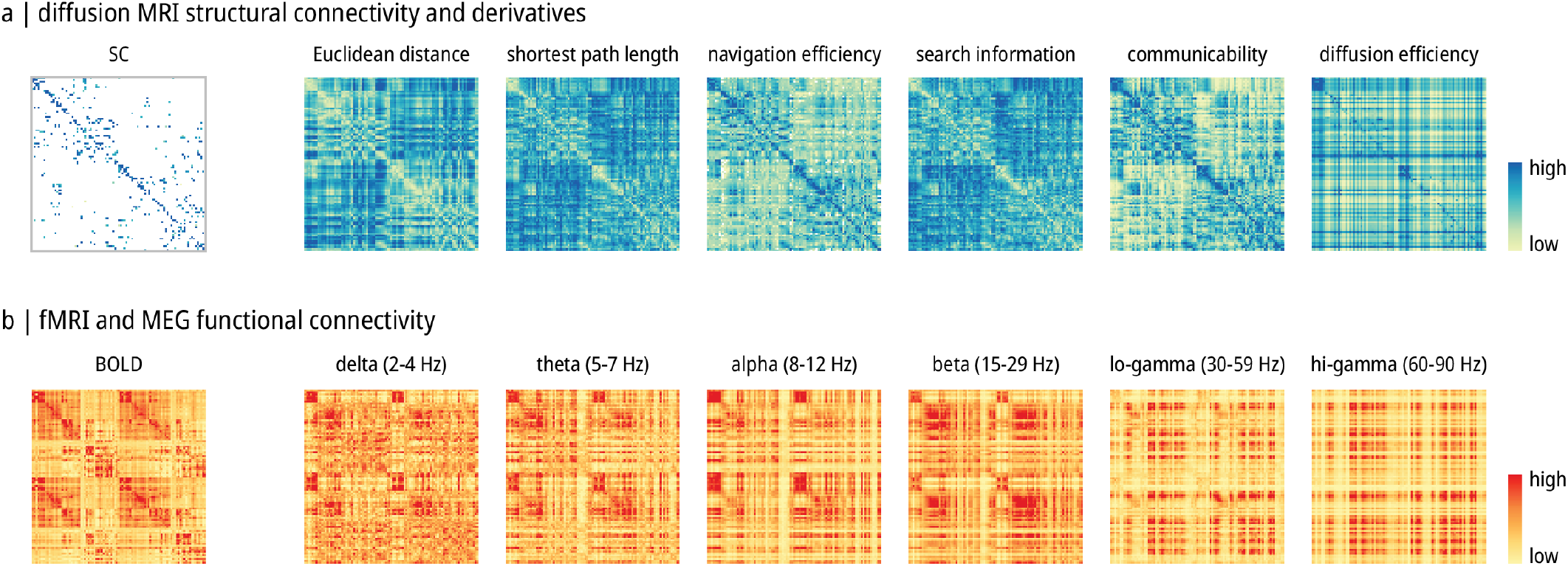
Structural and functional matrices. **(a)** Diffusion MRI structural connectivity (SC) and derived communication measures (Euclidean distance, shortest path length, navigation efficiency, search information, communicability, diffusion efficiency). Structural connectivity, navigation efficiency, communicability, diffusion efficiency matrices were log-transformed for better visualization. **(b)** Resting-state functional MRI (BOLD) connectivity and resting-state MEG functional connectivity in 5 canonical frequency bands (*δ, θ, α, β*, lo-*γ*, hi-*γ*). The colorbar covers the 2.5% to 97.5% percentile range if final values contain negatives, and 0 to 97.5% otherwise.

The functional connectivity matrices were derived in the same sample of participants using resting-state fMRI and MEG (Fig. 1b). FMRI functional connectivity was estimated using the conventional zero-lag correlation among regional BOLD time-series. MEG functional connectivity was estimated using amplitude envelope correlation (AEC) resolved in the six canonical electrophysiological frequency bands, including delta (*δ*; 2 to 4 Hz), theta (*θ*; 5 to 7 Hz), alpha (*α*; 8 to 12 Hz), beta (*β*; 15 to 29 Hz), low gamma (lo-*γ*; 30 to 59 Hz), and high gamma (hi-*γ*; 60 to 90Hz).

### Benchmarking local and global structure-function coupling

To comprehensively quantify structure-function coupling both locally and globally, we estimate structure-function coupling from four complementary perspectives. Global and local coupling describes the scale at which structure-function coupling is quantified (whole brain versus region-wise) [151]. Global coupling is estimated by constructing linear regressions relating the upper triangle of each SC predictor to the upper triangle of each FC outcome. The procedure generates a single *R*^2^ value, which is interpreted as the extent of structure-function coupling for the whole brain. Local coupling applies the calculation to each nodal connectivity profile, relating a region’s structural connectivity profile to its functional connectivity profile. This procedure generates a vector of coupling values corresponding to each brain region. To capture communication dynamics supported by structural connectivity, we also make a distinction between univariate and multivariate coupling [125, 151, 163]. In the multivariate case we take into account multiple communication measures estimated from the structural network, while in the univariate case we only use the structural connectivity matrix as the predictor (Fig. 2a).

**Figure 2.**
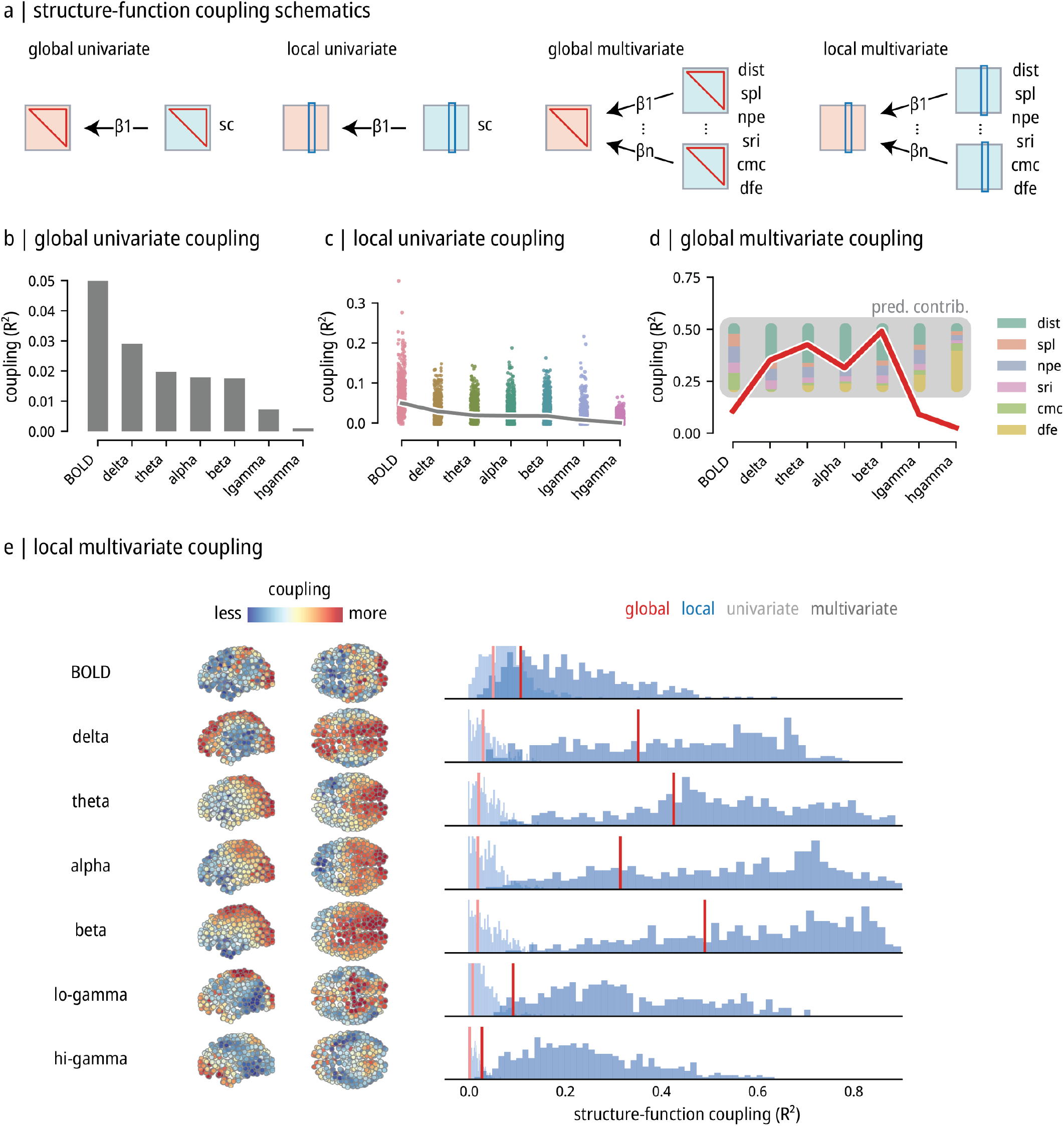
Local and global structure-function coupling. **(a)** Schematics of progressively detailed structure-function coupling measures. Structural connectivity matrix and its derivatives (shown as blue squares) are used to predict functional connectivity (shown as red squares) in a (multi)linear regression setting. For “global” coupling, the upper triangular matrix (shown in red triangles) is used. For “local” coupling, nodal profile (shown in blue rectangles) is used. **(b)** The global univariate coupling. Gray bars show the coupling value (adjusted *R*^2^) between structural connectivity and each type of functional connectivity. **(c)** The local univariate coupling. Scatters show structure-function coupling values for each node. Gray line shows the global univariate coupling from (b) for reference. **(d)** The global multivariate coupling. The red line shows structure-function coupling values using multiple predictors. Colored stacked bars show the ratio of predictor contributions calculated using dominance analysis for each regression. (e) The local multivariate coupling. Brain plots on the left show structure-function coupling values for each region and each functional connectivity type. Colorbar represents 2.5% to 97.5% percentile range. Distribution plots compare “global univariate” (lighter red line), “local univariate” (lighter blue bars), “global multivariate” (darker red line), and “local multivariate” (darker blue bars) settings. SC: structural connectivity, dist: Euclidean distance, spl: shortest path length, npe: navigation efficiency, sri: search information, cmc: communicability, dfe: diffusion efficiency.

Global univariate coupling – simply using structural connectivity as a predictor of functional connectivity – provides a baseline for characterizing structure-function coupling. Fig. 2b shows evidence of positive but overall weak coupling for all types of functional connectivity, with greatest values for BOLD-estimated and low-frequency MEG connectivity and smallest values for high-frequency MEG connectivity. By comparison, local univariate coupling – estimating structure-function coupling separately for each brain region – shows considerable regional heterogeneity (Fig. 2c). Notably, many regions show much greater structure-function coupling (shown by distributed points) compared to the corresponding global univariate coupling (shown by the gray line). Collectively, these results demonstrate that structure-function coupling exhibits considerable nuance and regional heterogeneity for both haemodynamic and electromagnetic networks.

We next consider global multivariate coupling, in which multiple regression is used to predict functional connectivity using a set of communication measure predictors derived from structural connectivity (Fig. 2d). Inclusion of multiple communication measures into the predictor set generally increases the estimated values of structure-function coupling, with adjusted *R*^2^ values ranging from 0.1 to 0.5. This increase potentially reflects the fact that different communication measures encode diverse types of dynamics that could be supported by the underlying structural connectivity matrix and therefore are better able to capture the emergent patterns of functional connectivity. Interestingly, we now observe the greatest structure-function coupling using MEG-estimated functional connectivity, particularly in the theta, alpha and beta rhythms (Fig. 2d; red line).

To provide greater detail about the contribution of each communication predictor, we additionally show a stacked bar plot depicting their percent dominance, a measure of each predictor’s importance (see *Methods*; [7, 26]). We note two important trends. First, the greatest contributor is usually Euclidean distance, consistent with numerous reports showing that the prevalence and strength of connectivity is greater among proximal neural elements and smaller among distal neural elements [16, 43, 66, 87, 88, 93, 100, 101, 109, 137]. Second, we find that faster frequency bands are generally better predicted using decentralized communications measures (e.g. diffusion efficiency; shown in yellow), while slower frequency bands and BOLD connectivity is generally better predicted by centralized communication measures (e.g. shortest path length; shown in orange). Collectively, these results suggest that different forms of haemodynamic and electromagnetic coupling may arise from distinct communication protocols in structural networks.

Finally, we consider local multivariate coupling – predicting a region’s functional connectivity profile from its communication profiles according to multiple communication measures (Fig. 2e). Regional distributions of *R*^2^ values are shown on the left for each type of functional connectivity. A histogram of *R*^2^ values (local multivariate, dark blue histogram) is shown on the right. For completeness, and to facilitate comparison, we also present results alongside other types of coupling (global univariate, light red line; global multivariate, dark red line; local univariate, light blue histogram). As expected, we observe overall greater structure-function coupling when using a local multivariate model, as it explicitly allows for both regional heterogeneity as well as multiplexed communication, with *R*^2^ often exceeding 0.6. Consistent with the results for the global multivariate model in Fig. 2d, we observe greatest structure-function coupling in the slower and intermediate MEG frequency bands, notably delta, theta, alpha and beta (Fig. 2e).

We finally look at the spatial distribution of structure-function coupling (Fig. 2e). BOLD coupling, similar to previous reports, delineates the canonical sensory-association axis, with greater coupling in sensory cortex and lower coupling in association cortex. By comparison, regional patterns of structure-function coupling for different MEG frequency bands tend to broadly follow a saggital gradient, separating anterior prefrontal cortex (low structure-function coupling) from posterior cortex, namely superior parietal and occipital cortex (high structure-function coupling). This pattern is most prominent in the theta, alpha and beta bands. The slower delta band, and the faster low and high gamma bands, show a slightly divergent pattern, in which the greater structure-function coupling is observed in medial superior cortex and weaker coupling elsewhere.

### Topological and intrinsic functional organization

In the previous section we systematically compared multiple coupling models. Here we examine in greater detail the “local multivariate” model as it takes into account both regional heterogeneity and multiple types of inter-regional signaling. We first examine whether regional differences in structure-function coupling are correlated with overall structural connectivity, as measured by the total weight of structural connections incident on a node (termed weighted degree; Fig. 3a; left). The logic is that regions with greater structural connectivity may be more prominently involved in inter-regional communication and therefore display greater structure-function coupling. Fig. 3a (right) shows the correlation between regional structural weighted degree and the extent of structure-function coupling. We observe moderate positive correlations for BOLD functional connectivity and delta band functional connectivity, but small and nonsignificant correlations for faster MEG frequency bands. This suggests that the background influence of total structural connectivity weight differently contributes to different frequency bands, and may be mainly expressed at slower time scales but not at faster time scales.

**Figure 3.**
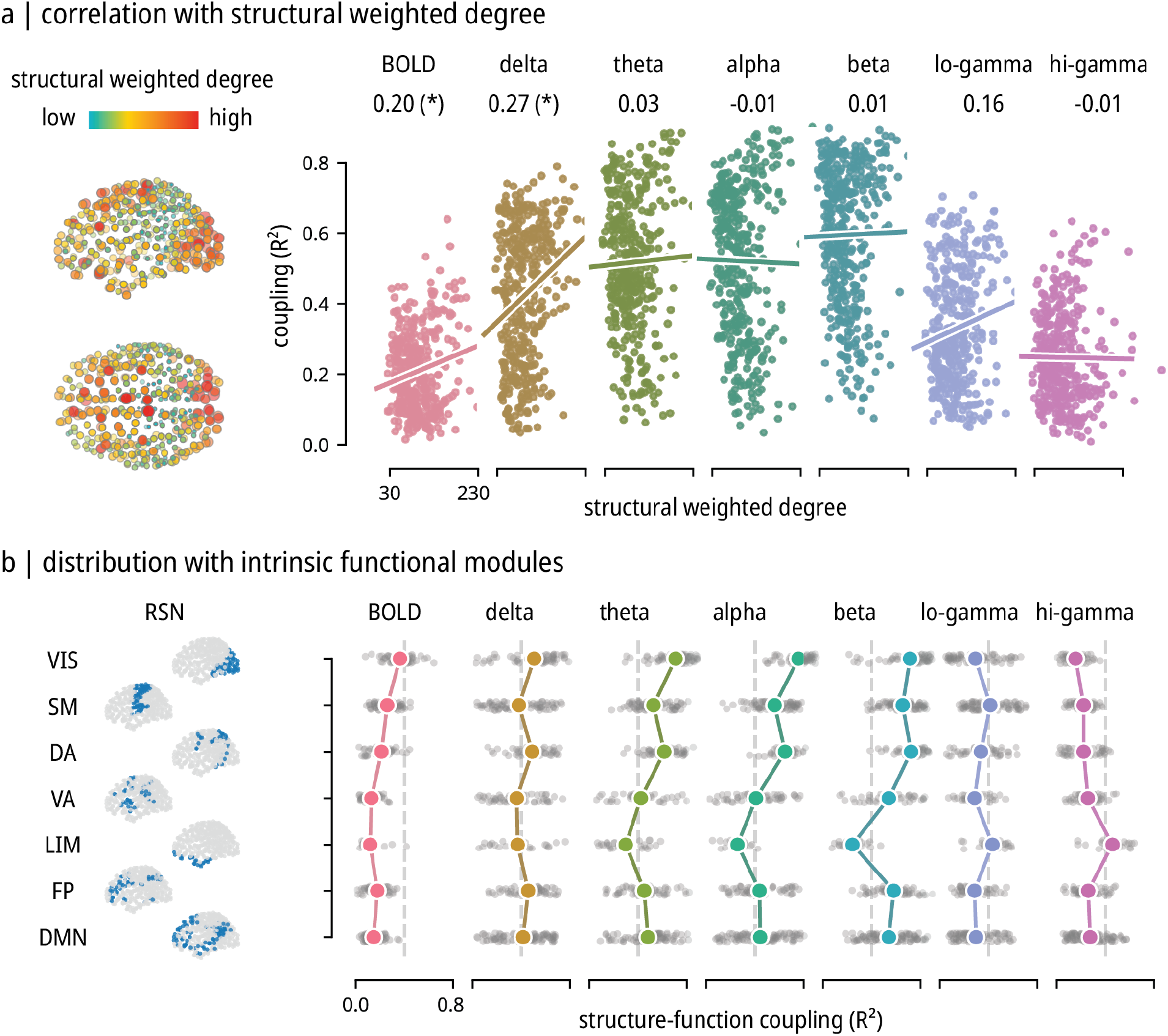
Topological and intrinsic functional organization. **(a)** Correlation between structure-function coupling and structural weighted degree. The structural weighted degree calculated as nodal average of the weighted structural connectivity matrix is shown on the left. Colorbar represents 2.5% to 97.5% percentile range. Its correlation with coupling values are shown on the right with scatters and regression lines. Pearson correlation coefficients are shown above each panel. Asterisks represent statistically significant correlations under the “spin null” (one-sided permutation test, *p <* 0.05). **(b)** Distribution of structure-function coupling in fMRI-derived intrinsic functional networks. Scatter brains on the left display the 7 networks from [161]. Gray scatters on the right show the coupling values belonging to each intrinsic network and colored points represent the mean value. Vertical dashed gray lines mark *R*^2^ = 0.4 on the x-axis. VIS: visual, SM: somatomotor, DA: dorsal attention, VA: ventral attention, LIM: limbic, FP: fronto-parietal, DMN: default mode network.

We next ask how well regional patterns of structure-function coupling align with the canonical intrinsic functional networks [119, 161] (Fig. 3b). These networks or modules are derived from resting state fMRI BOLD measurements and are thought to be the building blocks of higher cognition [146]. Specifically, we compute the mean structure-function coupling (estimated by *R*^2^) for all regions in a particular network. Consistent with previous reports, we find greater structure-function coupling in primary sensory and motor cortex (e.g. visual and somatomotor networks) and lower structure-function coupling in transmodal cortex (e.g. default network). This trend is most evident in BOLD and slower MEG frequency bands. In functional networks estimated using faster MEG rhythms, there are also deviations from this overall pattern; notably, the extent of structure-function coupling in the limbic network is sensitive to frequency band, with lower structure-function coupling in the beta band, and greater coupling in the low and high gamma bands. Importantly, the spatial organization of MEG structure-function coupling is reminiscent of but distinct from BOLD, and dependent on the rhythm being considered, which suggests potentially different modes of intrinsic functional organization in MEG [8, 115]. Collectively, these results show that structure-function coupling is spatially highly organized and potentially depends regional affiliation with macroscale functional gradients and/or regional micro-architecture. We explore this possibility in the next subsection.

### Structure-function coupling reflects cortical micro-architecture

Given that structure-function coupling is regionally specific and exhibits distinct topological and modular organization across frequency bands, we next seek to relate the coupling patterns to the hierarchical organization of the cortex. First, we apply the archetypal sensorimotor-association axis, a composite continuous ranking scale that combines multiple histological and imaging gradients [139] (Fig. 4a; left). Consistent with the intuition developed in the previous subsections, we generally observe negative correlations between the hierarchical position of a region and its structure-function coupling, indicating a gradual decoupling of structure and function as one moves from unimodal to transmodal cortex. This effect is particularly prominent in BOLD (*r* = − 0.63) and theta (*r* = − 0.55), alpha (*r* = − 0.67), and beta (*r* = − 0.57) MEG frequency bands. Interestingly, effect sizes gradually become smaller in higher frequencies and, although not statistically significant, the trend is reversed in the high gamma regime, potentially suggesting greater structure-function coupling in transmodal cortex.

**Figure 4.**
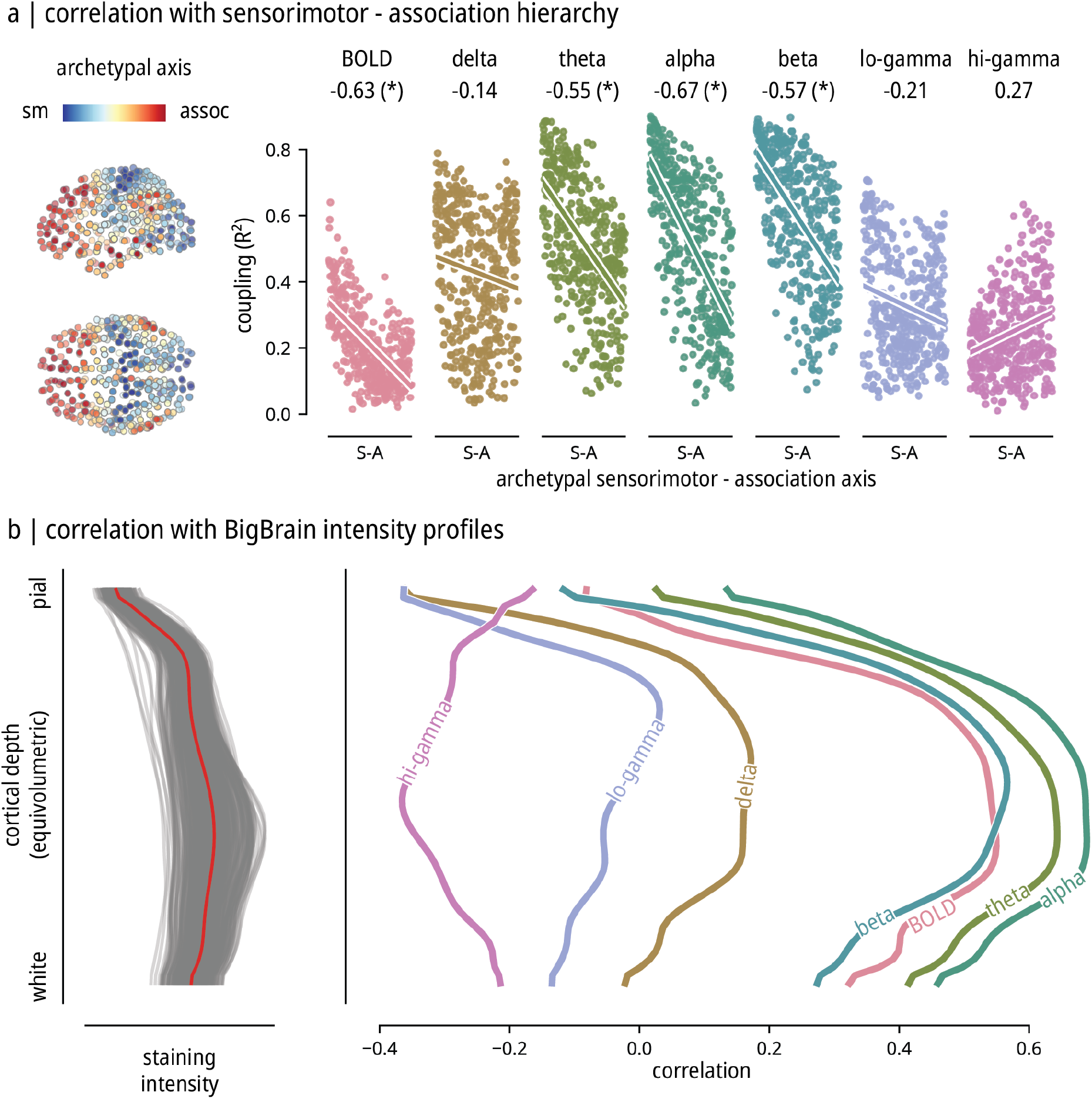
Hierarchical and cytoarchitectural organization. **(a)** Correlation between structure-function coupling and the sensorimotor-association hierarchy. The archetypal axis from [139] is shown on the left. The correlations with structure-function coupling are shown on the right with scatters and regression lines. Pearson correlation coefficients are shown above each panel. Asterisks represent statistically significant correlations under the “spin null” (one-sided permutation test, *p <* 0.05). **(b)** Correlation between structure-function coupling and BigBrain cytoarchitectural intensity profiles. The staining intensity values from pial to white surfaces are shown on the left, with gray lines representing different cortical regions, and red line representing the mean. The Pearson correlation between coupling values and staining intensity at each cortical depth are shown on the right. Lines correspond to functional connectivity types.

The previous result shows that structure-function coupling in both electromagnetic and haemodynamic networks depends on a region’s position in the cortical hierarchy. This raises the possibility that regional differences in structure-function coupling may reflect intrinsic differences in a more fundamental cortical feature, namely cytoarchitecture. Importantly, the laminar organization of the cortex determines the spatial organization of cell types, subcortical and cortico-cortical input and vascularization. As a result, numerous studies posit that different electromagnetic rhythms may potentially originate from different layers [105, 120, 121, 131]. We therefore asked whether regional differences in structure-function coupling could be explained by patterns of laminar differentiation.

To address this question, we used the 3D BigBrain histological atlas to estimate variations in cell density and size at multiple cortical depths [2, 103]. Merker cell-staining intensity profiles were sampled across 50 equivolumetric surfaces from the pial surface to the white matter surface to estimate laminar variation in neuronal density and soma size (Fig. 4b; left). We then correlate layer-specific intensity values (representing how prominent a particular layer is in a given region) with regional values of structure-function coupling (Fig. 4b; right). We generally observe the highest-magnitude correlations (positive or negative) in intermediate layers, corresponding to the granular layer IV [103], suggesting that regional differences in structure-function coupling mainly originate from this specific layer. The prominence of layer IV is positively correlated with structure-function coupling in BOLD and slower and intermediate MEG frequency bands (delta, theta, alpha and beta), potentially because layer IV receives many feedforward projections and has greater vascular density compared to other layers [39, 60, 123]. It is noteworthy that structure-function coupling in faster rhythms, particularly low gamma (30 to 59 Hz), is more prominent in more superficial layers. Collectively, these results highlight how structure-function coupling depends on laminar differentiation and cytoarchitectural gradients.

## DISCUSSION

The present report comprehensively quantifies patterns of structure-function coupling across the neocortex using both MEG and fMRI. We find four principal results. First, local models that allow for regional heterogeneity better capture structure-function relationships. Second, structure-function coupling is stronger in slower and intermediate frequency bands. Third, structure-function coupling in different bands is better captured by different communication models. Fourth, structure-function coupling is highly organized according to the sensorimotor-association axis and reflects patterns of laminar differentiation.

The nature of the structure-function relationship remains a key question in the field [138]. Multiple studies have investigated the relationship between anatomical connectivity and haemodynamic connectivity estimated from resting state fMRI at the global level [116, 138]. A parallel literature looks at how anatomical pathways support the emergence of neurophysiological oscillations and synchrony [29, 91, 115, 116, 134, 141]. In both cases, the focus has traditionally been on global structure-function coupling, under the assumption that there exists a single, consistent relationship between structural and functional connectivity across the brain.

We find that models that allow for regional heterogeneity offer greater nuance and anatomical detail for local structure-function coupling. In both BOLD functional connectivity networks and multiple MEG frequency bands, we find consistent patterns of greater structure-function coupling in unimodal cortex and lower coupling in transmodal cortex, tracing out the archetypal sensorimotor-association axis [61, 83, 139]. This organization potentially suggests that emergent patterns of inter-regional signaling gradually decouple from the underlying macroscale anatomical connectivity in trans-modal cortex. One hypothesis is that rapid evolutionary expansion of transmodal cortex “untethers” circuit configuration from the underlying transcriptomic gradients, resulting in a broader functional repertoire [25]. Patterns of structure-function coupling, which follow the archetypal sensorimotor-association axis, may therefore reflect underlying micro-architectural gradients.

Indeed, we find that regional patterns of structure-function coupling may potentially reflect patterns of cytoarchitecture. Namely, the magnitude of structure-function coupling in different frequency bands depends on regional differences in laminar differentiation, as estimated using the BigBrain histological atlas [2]. This is consistent with previous reports that have established laminar specificity for both neurophysiological rhythms and for neurovascular coupling. For instance, studies in both humans and animals have shown that in the visual cortex, alpha band activity is associated with deeper cortical layers like infragranular layers V to VI, while gamma band activity is associated with superficial cortical layers like supragranular layers I to III and granular layer IV [27, 59, 79, 80, 105, 121, 131, 135, 159]. Moreover, patterns of neurovascular coupling display a topography that is similar to what we find, with greatest vascularization density in layer IV [40, 120, 123]. At the same time, layer IV receives the majority of feedforward input [87, 89, 110, 152], consistent with our finding that regions with more prominent layer IV profile tend to display stronger coupling between macroscale structural and functional connectivity. How cortical micro-architecture shapes the relationship between macroscale structure and function remains an exciting question for the field.

The relationship between network configurations reconstructed from neurophysiology, fMRI and anatomical connectivity has received a lot of attention. For instance, a consistent finding is that fMRI network connectivity is mainly driven by slow oscillations [23, 37, 42, 45, 75, 82, 116, 127]. Structure-function coupling makes it possible to compare BOLD networks and MEG networks in a common, biologically-meaningful frame of reference. Indeed, we find that patterns of BOLD structure-function coupling typically resemble slower-frequency MEG structure-function coupling. At the same time, there are important inter-modality differences, as well as differences among the different MEG rhythms. Namely, while BOLD and delta structure-function coupling follows a unimodal-transmodal topography, structure-function coupling in intermediate bands (e.g. theta, alpha and beta) tends to follow a saggital gradient that resembles the dominant spectral power gradient [78, 127].

What could be driving the differences among structure-function coupling patterns in different oscillatory regimes? One possibility is that different rhythms entail different modes of signal exchange [11, 49, 71, 102]. By estimating structure-function coupling from the perspective of a spectrum of network communication protocols, we can infer what types of communication protocols contribute most to band-specific network configurations [5, 14, 124–126, 138, 162]. Here we find that centralized, routing-like protocols (e.g. shortest path length) and navigation-like protocols (e.g. distance-based navigation) better explain structure-function coupling in BOLD and slower-frequency MEG networks, whereas decentralized, diffusion-like protocols (e.g. random walk efficiency) better explain structure-function coupling in faster-frequency MEG networks. Collectively, these results suggest that the nature of signal exchange in neural circuits depends on the time scale, and that a spectrum of communication strategies may be implemented for different neurophysiological rhythms [4– 6, 51, 56, 94, 95].

The present results should be considered in light of two important methodological limitations. First, structural connectivity was estimated from diffusion MRI, a technique that is affected by systematic false positives and false negatives [21, 30, 35, 36, 57, 58, 69, 81, 99, 111, 118, 122, 132, 143, 160, 164]. We attempted to mitigate this issue by deriving a group consensus structural connectivity network that identifies consistent connections across many participants, but improvements in imaging technology and computational tractometry are still necessary. Second, many measures exist for quantifying correlations between functional time series [15, 22, 47, 50, 63, 77]. We used the conventional zero-lag Pearson correlation for BOLD and amplitude envelope correlation (AEC; [24]) for MEG because they are widely-used and most comparable to each other, and will therefore facilitate comparisons with other reports in the literature. Third, we adopted HCP-YA, which provides high-quality multimodal (MEG, fMRI and MEG) data in the same participants but is limited in sample size (*N* = 33). Future research on multimodal network comparisons could potentially either derive population-wide normative models using larger samples, or use precision imaging in more modalities with dense sampling in a smaller number of participants [97, 106]. For example, high-quality laminar-resolved functional data in parallel to diffusion acquisition is highly desired for further clarifying the origins and mechanism of how cortical rhythms arise from structure.

In summary, the present report comprehensively benchmarks patterns of structure-function coupling across multiple MEG frequency bands and BOLD functional connectivity. A consistent finding is that structurefunction coupling is not uniform but systematically organized across the brain, and parallels variations in cytoarchitecture. These results set the foundation for studying structure-function coupling as a phenotype of brain organization and open the door for multi-modal studies of structure-function relationships.

## METHODS

### Data preprocessing

Structural and functional data from 33 subjects (age range 22-35 years, no familial relationships) were obtained from the Human Connectome Project (HCP; s900 release [149]), including 3T structural MRI, multi-shell diffusion MRI, four resting-state functional MRI time series, and one resting-state MEG time series for each participant.

Briefly, MRI data were first preprocessed using HCP minimal preprocessing pipelines [53, 148]. Diffusion MRI scans were processed using MRtrix3 package [145]. Fiber orientation distributions were modelled using multi-shell multi-tissue constrained spherical deconvolution algorithm [38, 68]. White matter streamlines were then reconstructed using probabilistic tractography [144], and optimized using SIFT2 algorithm [133] to provide robust estimate of tract weights.

Resting-state fMRI scans (each approximately 15 minutes long, 4 scans for each participant) were corrected for gradient nonlinearity, head motion, and geometric distortions [153]. Further corrections include high-pass filtering (>2000s FWHM) for scanner drifts and ICA-FIX algorithm for additional noise [117, 153]. Both diffusion MRI and resting-state fMRI data were parcellated according to the Schaefer400x7 atlas [119]. Details of MRI data preprocessing are described in [104, 128].

Resting-state MEG scans (each about 6 minutes long) were processed with Brainstorm software [140], using a standard pipeline including notch filter, high-pass filter, bad channel removal, and automatic artifact removal (including heartbeats, eye blinks, saccades, muscle movements, and noisy segments). A linearly constrained minimum variance (LCMV) beamformer was used for source estimation on HCP fsLR4K surface. Parcellated time series were then estimated from the first principal component of the constituting sources’ time series according to the Schaefer400x7 atlas to facilitate comparisons with the MRI data [119]. Details of MEG data preprocessing are described in [127].

### Network reconstruction

Individual structural connectivity matrices were reconstructed with normalized streamline weights. A weighted consensus structural connectivity matrix was derived by identifying edges that consistently occur among multiple participants [17]. MRI functional connectivity edges were estimated as a Pearson correlation coefficient between time series for pairs of regions. A group-level consensus matrix was calculated by averaging the matrices across runs and subjects. MEG functional connectivity matrices were calculated using amplitude envelope correlation (AEC; [24]). Potential spatial leakage effects were corrected using an orthogonalization process [31] (see [127] for detailed discussion). AEC functional connectivity was derived for 6 canonical electrophysiological bands: delta (*δ*; 2 to 4 Hz), theta (*θ*; 5 to 7 Hz), alpha (*α*; 8 to 12 Hz), beta (*β*; 15 to 29 Hz), low gamma (lo-*γ*; 30 to 59 Hz), and high gamma (hi-*γ*; 60 to 90Hz).

### Network communication measures

Multiple network communication measures were computed on the sturctural connectivity matrix to estimate inter-regional communication dynamics across multiple putative communication protocols. They represent a spectrum of commonly-used measures ranging from centralized routing to decentralized diffusion. The network measures were implemented using the Brain Connectivity Toolbox ([114]; https://sites.google.com/site/bctnet), Brainconn Python Toolbox (https://github.com/FIU-Neuro/brainconn), and Netneurotools (https://github.com/netneurolab/netneurotools).

Before calculating the specific measures, Euclidean distance was taken between region centroids to represent the physical distance between regions. For some network measures, it is necessary to first define a connection length metric to quantify the cost of travelling through the edges. We used a monotonic weight-to-length transform adapted from [125] to derive the connection length matrix, which takes the form 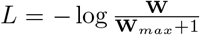. Applying the transform on the structural connectivity matrices effectively turns region pairs without direct structural connections to have infinite values in connection lengths, thus excluding such connections from the calculation of communication measures. For asymmetric measures, we take the mean value of the matrix and its transpose to make it a symmetric matrix [125]. We define the individual network communication measures as follows.

#### Shortest path length

Shortest path length denotes the shortest distance travelled between a source node and a target node [70, 74]. For weighted shortest path *π*_*s*→*t*_ = {*w*_*si*_, *w*_*jt*_ }, the shortest path lengths were calculated using Floyd-Warshall algorithm [46, 113, 158] on the connection length matrix *L*, and represented as 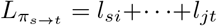.

#### Network navigation

Network navigation seeks to simulate a process where an agent or walker steps towards the neighbour node that is closest in distance to the target node [20, 96, 124, 126]. We used the Euclidean distance between nodes for navigation distance metric and derived navigation path lengths. For navigation between node *s* to *t*, if the walker successfully reached the target, the navigation path length is Λ_*st*_ = *d*_*si*_ + … + *d*_*jt*_, where *d*_*ij*_ is Euclidean distance. If the walker fails to reach the target, then Λ_*st*_ = ∞. Navigation efficiency is calculated as the inverse of navigation path length 1*/*Λ_*st*_.

#### Search information

Search information measures the amount of information necessary for a random walker to follow a given path, instead of taking other possible detours along the way [74, 112]. The metric was adapted to shortest paths on weighted networks in [55, 124]. For a random walker following the path *π*_*s*→*t*_ = {*w*_*si*_, *w*_*jt*_ }, the probability of staying on the path can be expressed as

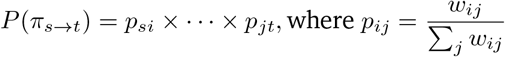

Search information can then be defined as

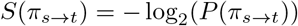

The group consensus structural connectivity matrix was used to simulate the random walk, and the connection length matrix was used to identify the shortest paths.

#### Communicability

Communicability between two nodes *i* and *j* is defined as the weighted sum of all paths and walks between those nodes [44]. For a weighted adjacency matrix *W*, communicability is calculated as

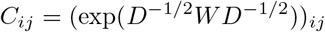

where 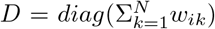 is the diagonal matrix of the generalized node degree matrix [3, 32, 151].

#### Diffusion efficiency

Diffusion efficiency is calculated as inverse of the mean first passage time, which itself is defined as the expected time steps for a random walker to reach the target node from the source node [54, 98]. Modelled as a Markov chain, the mean first passage time can be derived as follows. Let *P* be the transition matrix defined as *D*^−1^*W*, where *W* is the weighted adjacency matrix and *D* is the diagonal weighted degree matrix. Let *ω*_*i*_ be the probability vector corresponding to the stationary solution of the Markov process, and Ω as the column-wise probability vector matrix containing *ω*_*i*_. The fundamental matrix *Z* is computed as *Z* = (*I* − *P* +Ω)^−1^, where *I* is the identity matrix. The ergodicity property of a Markov chain for an undirected connected graph allows computing the mean first passage time as

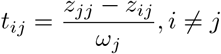

Diffusion efficiency is then calculated as 1*/t*_*ij*_. The group consensus structural connectivity matrix was used to simulate the random walk.

### Quantifying structure-function coupling

To estimate structure-function coupling, we used a set of linear regression models. The simplest form (“global univariate”) is

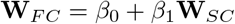

where **W**_*F C*_ denotes a consensus functional connectivity matrix from BOLD or MEG frequency bands, **W**_*SC*_ denotes a consensus structural connectivity matrix from diffusion MRI, and the regression is computed by taking the upper triangular values from each matrix. The extent of structure-function coupling is evaluated using the adjusted *R*^2^, the goodness of fit statistic.

Taking this notion further to the level of individual brain regions (“local univariate”), we compute a regression for the structural and functional connectivity profile of each node

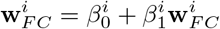

where **w**^*i*^ denotes connection profile for node *i* (i.e. the i^th^ column of the matrix). Diagonal elements were removed during the regression.

Another way to improve the “global univariate” model is to incorporate predictors derived from network communication models to account for potentially more complex processes of communication happening on the structural network, which we term “global multivariate”. The model is implemented as

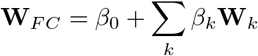

where *k* denotes predictor matrices: Euclidean distance, shortest path length, navigation efficiency, search information, communicability, and diffusion efficiency.

The final and the most complete form (“local multivariate”) is

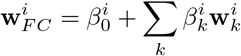

Similarly, *i* enumerates nodes and *k* enumerates predictor matrices.

### Cortical annotations

We used several common cortical annotations to contextualize the structure-function coupling patterns. Intrinsic functional networks are defined in [161] using resting-state functional MRI. Archetypal sensorimotor-association axis were defined in [139], approximating the putative sensory-fugal cortical hierarchy [83, 92] by fusing the ranks of 10 previously released brain maps in different modalities. BigBrain intensity profiles were generated using BigBrainWarp toolbox [2, 103], which measure the cytoarchitectural feature of cell staining intensity in 50 equivolumetric surfaces [154–156].

### Null model

To robustly estimate the statistical significance of correlation between nodal coupling and cortical annotations, we used spatial autocorrelation-preserving permutation null models, termed “spin tests” [1, 86, 150]. In this type of permutation test, the cortical surface is projected to a sphere and randomly rotated, generating permuted surface maps with preserved spatial autocorrelation. Using the spherical projection of the *fsaverage* surface for Schaefer400x7 atlas, the spherical coordinates of the parcels were defined by selecting the vertex closest to the center-of-mass of each parcel. Randomly sampled rotations were then applied on the sphere, and parcel values were reassigned based on the closest resulting parcel. Rotations were applied for one hemisphere and mirrored to the other. We generated 10,000 spin permutations (“Vázquez-Rodríguez” method) using *netneurotools* package (https://github.com/netneurolab/netneurotools) [151]. Details of spatially-constrained null models in neuroimaging were described in [86] (https://github.com/netneurolab/markello_spatialnulls) and are implemented in *neuromaps* package [84, 85]. This particular implementation of the spin test (“Vázquez-Rodríguez” method) was chosen based on the benchmarking results reported in [86], which showed that the method (a) was consistently the most conservative method in both simulations and empirical analyses, and (b) it was designed specifically for parcellated data and does not require discarding permutations when parcels are rotated into the medial wall.

### Predictor contributions

Contribution of network communication predictors in Fig. 2d was estimated using dominance analysis [7, 26], one of the procedures for interpreting multilinear regression models. It can account for multicollinearity and is sensitive to potential patterns in the model [72]. We adopted “total dominance” statistic for each predictor, which quantifies its contribution to the goodness of fit of the full model. It is estimated as the relative importance of each predictor by re-fitting all subset combinations of existing predictors. This function is implemented in *netneurotools* (https://github.com/netneurolab/netneurotools), which is adapted from the Dominance-Analysis (https://github.com/dominance-analysis/dominance-analysis) package.

## ACKNOWLEDGMENTS

We thank Justine Hansen, Laura Suárez, Vincent Bazinet, Filip Milisav, and Andrea Luppi for comments and suggestions on the manuscript. BM acknowledges support from the Natural Sciences and Engineering Research Council of Canada (NSERC), Canadian Institutes of Health Research (CIHR), Brain Canada Foundation Future Leaders Fund, the Canada Research Chairs Program, the Michael J. Fox Foundation, and the Healthy Brains for Healthy Lives initiative. ZQL acknowledges support from the Fonds de Recherche du Québec – Nature et Technologies (FRQNT).

## Notes

### Competing Interest Statement

The authors have declared no competing interest.

